# Pathogens spread by high-altitude windborne mosquitoes

**DOI:** 10.1101/2024.12.26.630351

**Authors:** R Bamou, A Dao, AS Yaro, C Kouam, K Ergunay, BP Bourke, M Diallo, ZL Sanogo, D Samake, YA Afrane, AR Mohammed, CM Owusu-Asenso, G Akosah-Brempong, CM Pambit-Zong, BJ Krajacich, R Faiman, MA Pacheco, AA Escalante, SC Weaver, R Nartey, JW Chapman, DR Reynolds, Y-M Linton, T Lehmann

**Affiliations:** Laboratory of Malaria and Vector Research, NIAID, NIH. Rockville, MD, USA; Malaria Research and Training Center (MRTC) / Faculty of Medicine, Pharmacy and Odonto-stomatology, Bamako, Mali; Walter Reed Biosystematics Unit (WRBU), Smithsonian Institution Museum Support Center, Suitland Maryland, USA; Department of Entomology, Smithsonian Institution, National Museum of Natural History, Washington DC, USA; One Health Branch, Walter Reed Army Institute of Research, Silver Spring, MD, USA; Department of Medical Microbiology, University of Ghana Medical School, University of Ghana.; Department of Animal Biology and Conservation Science, University of Ghana.; Biotechnology and Nuclear Agriculture Research Institute, Ghana Atomic Energy Commission, 25 Accra Ghana; Biology Department/Institute of Genomics and Evolutionary Medicine (iGEM), Temple University, Philadelphia, PA, USA; Department of Microbiology & Immunology and World Reference Center for Emerging Viruses and Arboviruses, University of Texas Medical Branch, Galveston, Texas, USA; Centre for Ecology and Conservation, and Environment and Sustainability Inst., University of Exeter, Penryn, Cornwall, UK; Department of Entomology, College of Plant Protection, Nanjing Agricultural University, Nanjing, P. R. China; Natural Resources Institute, University of Greenwich, Chatham, Kent, UK; Rothamsted Research, Harpenden, Hertfordshire, Kent, UK

**Author notes:** Correspondence: Tovi Lehmann.

**Keywords:** *Aedes*, *Anopheles*, arbovirus, *Culex*, disease-spread, filariae, dispersal, high-altitude windborne migration, malaria, mosquito-borne pathogen, *Plasmodium*, surveillance

## Abstract

Recent studies have revealed that many mosquito species regularly engage in high-altitude windborne migration, but its epidemiological significance was debated. The hypothesis that high-altitude mosquitoes spread pathogens over large distances has not been directly tested. Here, we report for the first time that high-altitude windborne mosquitoes are commonly infected with arboviruses, protozoans, and helminths affecting vertebrates and humans, and provide the first description of this pathogen-vector aerial network. A total of 1,017 female mosquitoes (81.4%, N=1,249) intercepted on nets suspended from helium balloons at altitudes of 120-290 m above ground over Mali and Ghana were screened for infection with arboviruses, plasmodia, and filariae, using pan-genus qPCR analyses followed by sequencing of positive samples. The mosquito fauna collected at altitude comprised 61 species, across 9 genera, dominated by *Culex*, *Aedes,* and *Anopheles*. Infection and infectiousness rates of high-altitude migrant mosquitoes were 7.2% and 4.4% with plasmodia, 1.6% and 0.6% with filariae, 3.5% and 1.1% with flaviviruses, respectively. Nineteen mosquito-borne pathogens were identified, including three arboviruses: dengue, West Nile and M’Poko viruses, 13 putative plasmodia species including *Plasmodium matutinum* and *P. relictum*, three filariids, including *Pelecitus* spp., 27 insect-specific viruses and 5 non-mosquito-borne pathogens (e.g., *Trypanosoma theileri*). Confirmed head-thorax (disseminated) infections of multiple pathogens in multiple mosquito species, eg., *Culex perexiguus*, *Coquilletidia metallica*, *Mansonia uniformis*, and *Anopheles squamosus* provides evidence that pathogens carried by high-altitude windborne mosquitoes are infectious and likely capable of infecting naïve hosts far from their starting location. This traffic of sylvatic pathogens may be key to their maintenance among foci as well as initiating outbreaks away from them.

## Introduction

Windborne insect migration at altitude occurs regularly on massive scales in terms of biomass and distance that can extend up to hundreds of kilometers per night (Reynolds et al. 2006, Hu et al. 2016, Florio et al. 2020, Huang et al. 2024). Migration, defined as a persistent movement temporarily unaffected by immediate cues for food, reproduction, or shelter, with a high probability of relocating the animal to a new environment (Dingle and Drake 2007, Chapman et al. 2015) fits these flights and will be used here. Insect disease vectors, pests, and species vital for ecosystem vigor are common among high-altitude flyers (Pedgley et al. 1995, Reynolds et al. 1996b, Chapman et al. 2002, Reynolds et al. 2006, Chapman et al. 2015, Hu et al. 2019, Huestis et al. 2019, Wotton et al. 2019, Yaro et al. 2022, Huang et al. 2024). We poorly understand these movements in tropical mosquitoes and how they affect survival and reproductive success. Questions about their impacts on mosquito range expansion, inter-continental invasion, spread of insecticide resistance, and spread of vector-borne diseases have dominated the field (Kay and Farrow 2000, Eagles et al. 2013, Sedda and Rogers 2013, Surendran et al. 2020, Beeton et al. 2022, Lehmann et al. 2023b, Lehmann et al. 2023a). The hypothesis of pathogens spread by high-altitude windborne mosquitoes is not new (Garrett-Jones 1950, 1962, Sellers et al. 1977, 1982, Sellers 1989, Ming et al. 1993, Pedgley et al. 1995, Kay and Farrow 2000, Reynolds et al. 2006), but was supported largely by epidemiological and meteorological inferences, while direct evidence of the regularity of such movements, particularly of the infection of high-altitude windborne mosquitoes has been elusive. Recent studies in Africa revealed that many mosquito species engage in windborne migration at altitude, i.e., 40—290 m above ground level (agl) on a regular basis (Huestis et al. 2019, Yaro et al. 2022, Atieli et al. 2023) involving myriads of other insects (Drake and Reynolds 2012, Hu et al. 2016, Florio et al. 2020). Additional support for this hypothesis was provided by the findings that, among the migrants, approximately 90% of the mosquito females (mostly gravid) had had previous exposure to vertebrate blood, that the flights coincided with the disease-transmission season, and many of these species have been previously implicated as vectors of pathogens (Huestis et al. 2019, Yaro et al. 2022, Lehmann et al. 2023a). Here, we show migrant mosquitoes at altitude have high rates of infection with arboviruses, plasmodia, and filariae. Furthermore, migrating mosquitoes are not only infected i.e., exposed to these pathogens, but are already likely infectious, i.e., presenting a disseminated infection to the haemocoel and likely to the salivary glands, highlighting their capacity to transmit pathogens upon landing in a new territory.

## Results

### Aerial mosquito diversity

Of 1,249 mosquitoes collected at altitude (120–290 m agl) over West Africa during 191 collection nights between 2018 and 2020 and subjected to molecular analysis, the specific identity of 782 was confirmed by mitochondrial *COI* barcode sequencing (468 were identified to subfamily; Table S1), yielding 60 species across 9 genera (Fig. 1, Table S1). Diversity of *Culex* was highest (26 species), followed by *Aedes* (11), *Anopheles* (10), *Coquilittidia* (3), *Uranotaenia* (3), *Mansonia* (3), *Mimomyia* (2), and *Eratmopodites* (1), *Lutzia* (1). Among those identified to species, *Cx. watti* and *Cx. perexiguus* were the dominant taxa, together comprising half of the collections, followed by *Cx. cf. watti MAFP5.C5, Cq. metallica*, *Cx. univitattus*, and *Cx. neavei*; altogether comprising 69% of the collection (Fig. 1a). The next group of moderately common taxa included six species, e.g., *Ae. argenteopunctatus, An. squamosus, Ae. quasiunivitattus, Cx. antennatus*, and *Cx. nebulosus*, each represented by 2–3% of the specimens, followed by 21 species rare in our collection, e.g., *Ma. africana,* and *An. gambiae* s.s., each represented by ∼0.2% (1–2 specimens/species; Figs. 1 and S1). Identified taxa included primary vectors of malaria (*An. gambiae* s.s., *An. coluzzii*) and arboviruses, e.g., West Nile virus (*Cx. univitattus*), and Rift Valley fever virus (*Ae. mcintoshi*) (Diallo et al. 2005, Tandina et al. 2018, Diallo et al. 2019, Lehmann et al. 2023a). Females comprised 85% of identified specimens, with no evidence for inter-species heterogeneity in the sex composition (P=0.06, Table S2). Overall, gravid females consisted of 44% of identified species with fractions varying between 28% (*Cx. univitattus*, N=25) to 80% (*Cx. perexiguus*, N=123, among-species heterogeneity P=0.001, Table S3).

**Fig. 1.**
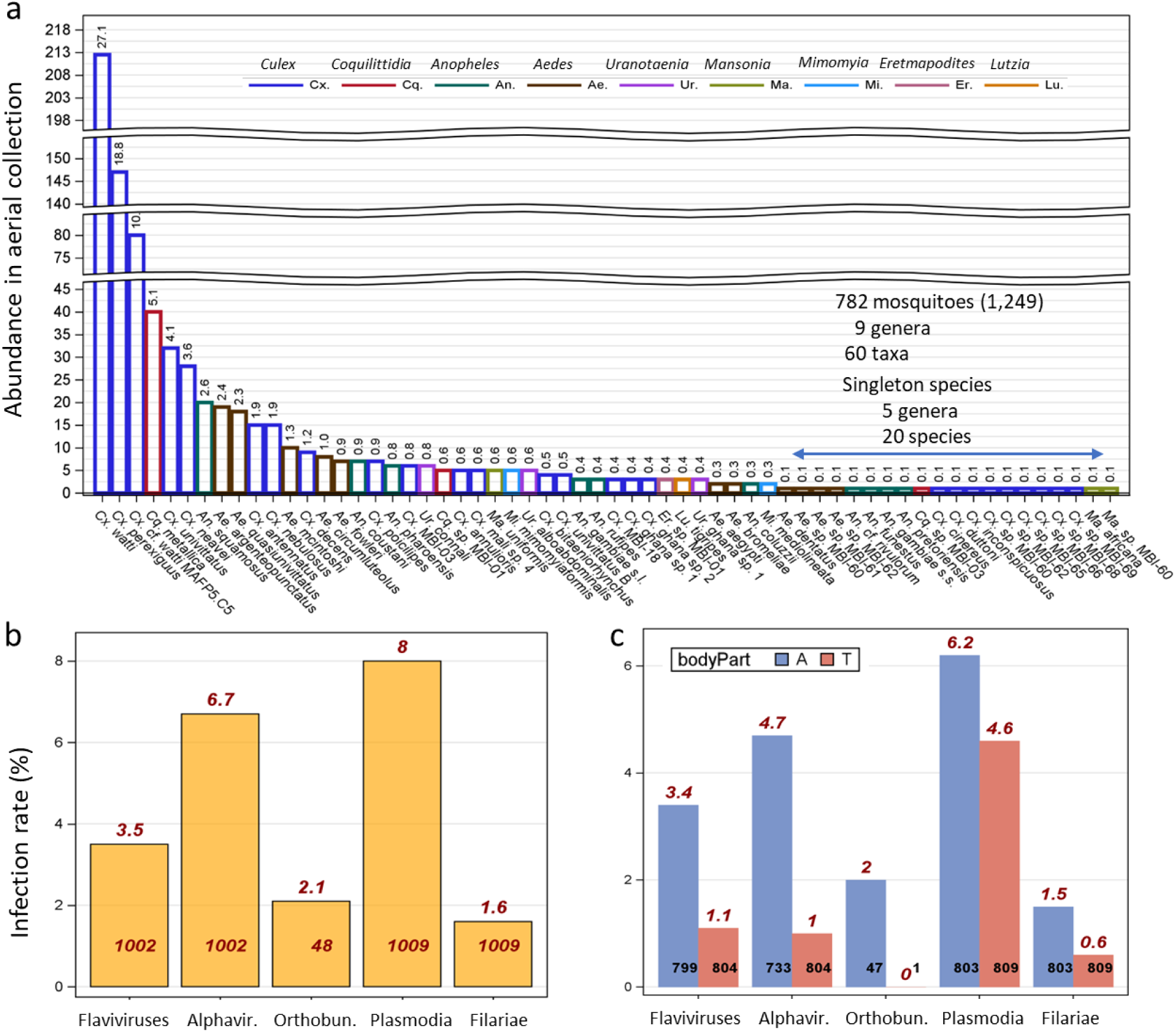
Mosquito species composition at altitude and infection rates with major pathogen groups. a) Mosquitoes identified to species are shown (N=782, note: breaks in the Y-axis; number of specimens per species). The percentage of the total specimens per species is shown above bars. Species represented by a single specimen are grouped under the blue line. b) Overall infection rates (per mosquito) of high-altitude windborne mosquitoes with select pathogen groups. (c) Infection rate in abdominal tissues (A, blue) and in head-thorax tissues (T, red). Infection rates expressed as percentages are shown above bars (red) and corresponding sample sizes are shown at the base.

### Aerial pathogen diversity

Pan-genus qPCR/PCR detection assays targeting Plasmodia, filariae, flaviviruses and alphaviruses were performed on 1,009 female mosquitoes that were captured at altitude, consisting of 803 abdomens, 809 thoraces, and 194 whole-body specimens. Overall mosquito infection rates (infection of any mosquito part) with each pathogen group varied between 1.6% (filariae) and 8.0% (plasmodia; Fig. 1b, Table 1, S4). Infection rates for flaviviruses, plasmodia, and filariae were higher in abdominal tissues than in head-thorax tissues (Fig. 1b, Tables 1 and S1), probably reflecting residual pathogen DNA/RNA after exposure in bloodmeal (see also Table S4) and early infection before pathogen dissemination beyond the midgut (abdomen) during the short life span of adult mosquitoes (Gillies and Wilkes 1965, Charlwood et al. 2012). Sequence comparisons confirmed pathogen infections in high-altitude mosquitoes with all pathogen groups except alphaviruses; hence, alphaviruses were excluded from subsequent analyses. Metagenomic sequences of 48 samples found positive for pan-flavivirus, pan-filaria and pan-plasmodium infection revealed infection with viruses of other families, as well as with non-mosquito-borne pathogens (Tables 1 and S4). A total of 20 vertebrate-mosquito pathogens were detected in this modest sample, including two flaviviruses (West Nile virus, dengue virus), one orthobunyavirus (M’Poko virus), 13 avian plasmodium species (>=3% *cytb* gene sequence dissimilarity, see Methods), and four filariid nematodes (Table 1). The Plasmodia included the cosmopolitan *P. matutinum*, *P. relictum*, and *P. vaughani* and the African-endemic *P.* sp. MALNI02 (MalAvi lineage) previously detected in Blue-billed Malimbe, *Malimbus nitens*, in Gabon (Table 1, [Malavi]). The three filarial taxa, members of different genera included *Cardiofilaria sp.*, *Pelecitus sp.* as well as a taxon related to or member of the genus *Loa* (Table 1). Natural hosts of these taxa include both birds and mammals (Table 1). These pathogens represent sylvatic and zoonotic species, e.g., WNV, which are transmitted by mosquitoes mainly among birds (but see DENV, Table 1). Several insect-specific viruses (viruses that infect insects but are incapable of infecting vertebrates, e.g., Barkedji virus and Nienokue virus), as well as non-mosquito-borne pathogens, e.g., *Trypanosoma theileri* and *Haemoproteus coraciae* were also detected (Table S4). Co-infection between pairs of pathogen groups (flaviviruses, plasmodia, and filariid nematodes) did not depart from random expectations when tested in whole body, abdomens, and head-thorax body parts (P>0.2, Fisher exact tests, not shown).

**Table 1.** Overall infection rates in female mosquitoes intercepted at altitude (120-290 m above ground) with select pathogens.

| Pathogen | Overall<sup>a</sup> (N) | Abdomen (N) | Head-thorax (N) | Method | Main natural hosts | Human Impact<sup>f</sup> |
| --- | --- | --- | --- | --- | --- | --- |
| **Flaviviruses** | 3.5% (35/1,002) | 3.4% (27/799) | 1.1% (9/804) | Pan-Flavivirus RT-qPCR | Vertebrates/Arthropod | Human, livestock[Z,P] |
| - West Nile Virus (lineage 1a)<sup>b</sup> | 0.2% (2/1,002) | 0.25% (2/799) | 0% (804) | Sanger | Birds/Mosquito | Human, horse [Z] |
| - dengue (lineage 2)<sup>c</sup> | 0.2% (2/1,002) | 0.25% (2/799) | 0% (804) | Sanger | Primates/mosquitoes | Human, primates [P] |
| **Peribunyaviridae; Orthobunyavirus** | 2.1% (1/48) | 2.1% (1/47) | 0% (1) | Metagenomics | Vertebrates/Arthropod | Human, livestock [Z] |
| - M'Poko virus<sup>d</sup> | 2.1% (1/48) | 2.1% (1/47) | 0% (1) | Metagenomics | Birds/Mosquito | Human [Z] |
| **Alphaviruses<sup>e</sup>** | 6.7% (67/1,002) | 1.0% (8/799) | 4.7% (38/804) | Pan-Alphavirus RT-qPCR | Vertebrates/Arthropod | Human, livestock[Z] |
| **Plasmodia** | **8.0% (81/1,009)** | **6.2% (50/803)** | **4.6% (37/809)** | Pan-Plasmodium qPCR | Birds/Arthropod | Human[P,Z] |
| - *P. AFR006* | 0.1% (1/1009) | 0% (0/803) | 0.1% (1/809) | Sanger | Birds/Mosquitoes | none/unknown |
| - *P. AFR146* | 0.4% (4/1009) | 0.25% (2/803) | 0.5% (4/809) | Sanger | Birds/Mosquitoes | none/unknown |
| - *P. AUTINF03* | 0.2% (2/1009) | 0.25% (2/803) | 0% (0/809) | Sanger | Birds/Mosquitoes | none/unknown |
| - *P. CAMBRA02* | 0.1% (1/1009) | 0.1% (1/803) | 0% (0/809) | Sanger | Birds/Mosquitoes | none/unknown |
| - *P. TCHSEN01* | 0.2% (2/1009) | 0.25% (2/803) | 0.1% (1/809) | Sanger | Birds/Mosquitoes | none/unknown |
| - *P. CXPER01* | 1.2% (12/1009) | 0.7% (6/803) | 0.7% (6/809) | Sanger | Birds/Mosquitoes | none/unknown |
| - *P. matutinum* | 1.3% (13/1009) | 0.5% (4/803) | 0.9% (7/809) | Sanger | Birds/Mosquitoes | none/unknown |
| - *P. relictum* | 0.6% (6/1009) | 0.5% (4/803) | 0.3% (2/809) | Sanger | Birds/Mosquitoes | none/unknown |
| - *P. vaughani* | 0.7% (7/1009) | 0.5% (4/803) | 0.1% (1/809) | Sanger | Birds/Mosquitoes | none/unknown |
| - *P. MALNI02* | 0.1% (1/1009) | 0.1% (1/803) | 0% (0/809) | Sanger | Birds/Mosquitoes | none/unknown |
| - *P. mali sp. 1* | 0.8% (8/1009) | 0.25% (2/803) | 0.1% (1/809) | Sanger | Birds/Mosquitoes | none/unknown |
| - *P. mali sp. 2* | 0.6% (6/1009) | 0.5% (4/803) | 0.1% (1/809) | Sanger | Birds/Mosquitoes | none/unknown |
| - *P. ghana sp. 1* | 0.4% (4/1009) | 0.25% (2/803) | 0.3% (2/809) | Sanger | Birds/Mosquitoes | none/unknown |
| **Filariae (Onchocercidae)** | **1.6% (16/1,009)** | **1.5% (12/803)** | **0.6% (5/809)** | Pan-Filariae qPCR | Vertebrates/Arthropod | Human, livestock |
| - *Cardiofilaria sp.* | 0.1% (1/1009) | 0.1% (1/803) | 0% (0/809) | Sanger | Birds/Mosquitoes | none/unknown |
| - *Loa like sp.* | 0.1% (1/1009) | 0.1% (1/803) | 0% (0/809) | Sanger | Unknown | none/unknown |
| - *Pelecitus sp.* | 0.1% (1/1009) | 0.1% (1/803) | 0% (0/809) | Metagenomics | Birds/Mammals | Human[Z] |

<sup>a</sup> Mosquito infection rate (regardless of bodypart, including whole-body mosquitoes) using PCR and metagenomics (positive number/number tested)
<sup>b</sup> One mosquito infection confirmed by Sanger (lineage 1a); other infection confirmed by WNV-specific qPCR (not sequencing or metagenomics)
<sup>c</sup> Infection was detected in one mosquito abdomen and another whole body female
<sup>d</sup> Infection was detected and confirmed by metagenomics after it was detected as positive to flaviviruses
<sup>e</sup> Infection with alphaviruses could not be confirmed by Sanger sequencing (N=67) nor by NGS metagenomics (N=48).
<sup>f</sup> Direct effect on human health or indirect by affecting health of livestock [P and Z denote primary and zoonotic host, respectively]

### Pathogen-mosquito relationships

Infection with plasmodia, filariae or flaviviruses was detected in 21 mosquito species intercepted at altitude, with an overall infection rate of 12.7% per female (N=1,009), 10.7% in abdomens (N=803), and 6.3% in head-thorax sections (N=809). These rates varied among mosquito species, but no systematic relationship between species’ sample size and pathogen prevalence were observed (Figures 2a and S2). Both *Uranotaenia connali* and *Cx. perexiguus* exhibited significantly higher overall infection rates than the means for all mosquito species (P<0.05, 2 sided exact binomial tests (Figures 2a and S2). Importantly, 15 mosquito species exhibited disseminated (head-thorax) infections with these pathogen groups, conditions required for infection of salivary glands and transmission competence (Fig. 2b and S3). Positive relationships between abdominal and head-thorax infection shown by high values in both the Y and X axes is driven by susceptible, competent vectors that are more commonly exposed by preferably feeding on natural host species of these pathogens (Fig. 2b). *Ur. connali*, *Ma. uniformis*, *Cq. metallica*, and *Ae. circumluteolus* are such species (Fig. 2b). *Culex perexiguus* is mapped close to the regression line, yet with 9% head-thorax infection rate and 17% abdominal infection rate, is flagged as a vector of potential importance, an inference that is compounded by its high abundance at altitude (Figs. 1a and 2b).

**Figure 2.**
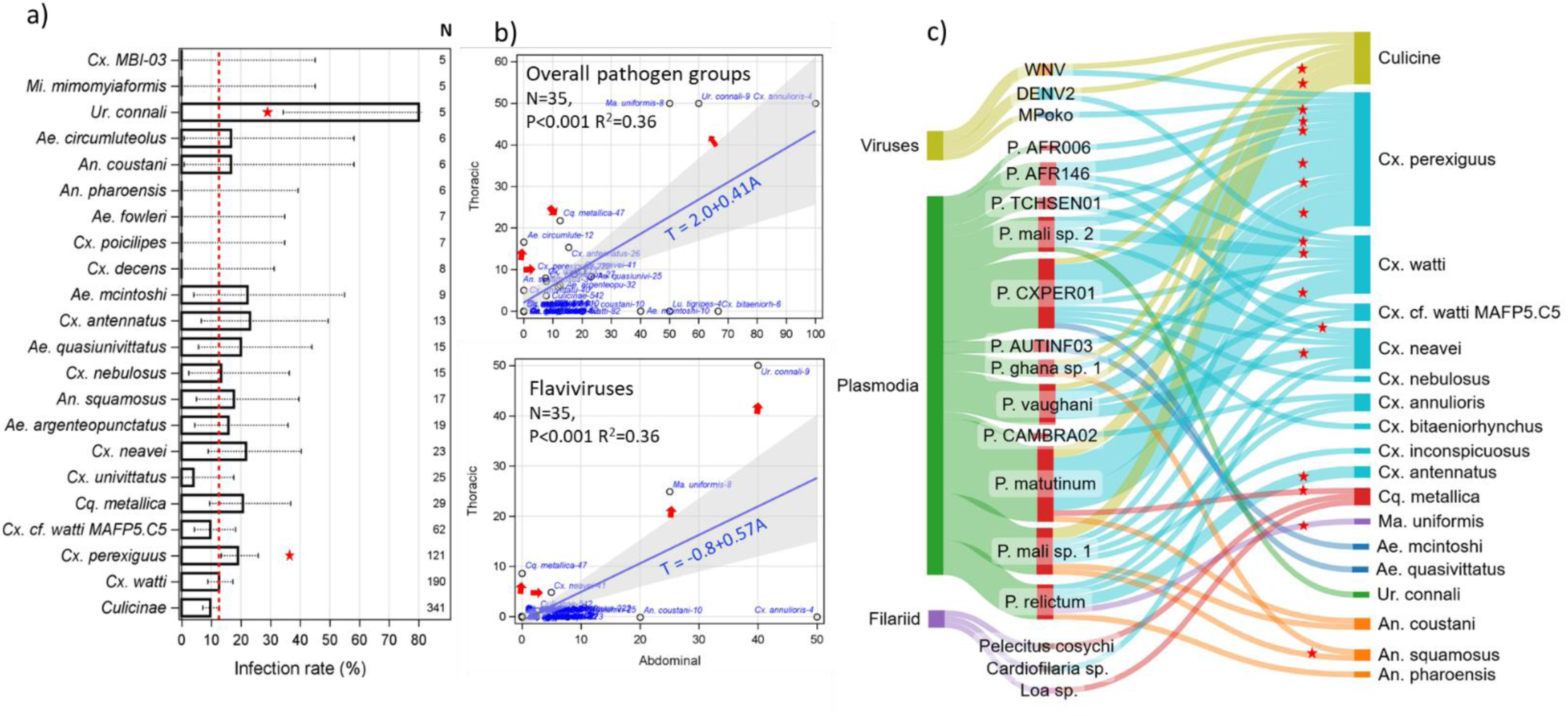
High-altitude overall mosquito infection rate per species (a) in head-thorax vs. abdominal tissues (b), and with specific vertebrate pathogens (c). a) Overall infection rates per mosquito species (N≥5, above bars) based on pan-genus PCR-based assays for flaviviruses, plasmodia, and filariae with 90% CI. Higher infection rate (P<0.05, 2 side Binomial test) than the rate across all mosquito species (12.7%, red line) are indicated by stars. b) Relationship between disseminated (head-thorax) infection and exposure (abdominal) infection by species (abbreviated species name is followed by their sample size), with linear regression weighted by sample size (blue) and 95% CI as reference to identify outliers (red arrows). Infection with all pathogen groups (top) and with flaviviruses (bottom) are shown (see Fig. S5 for plasmodia and filariae). c) Sankey diagram showing mosquito species infection with pathogens confirmed by sequencing. Connective band is proportioned to infection rate. Red stars indicate disseminated infection.

Sequencing from positive mosquitoes revealed 19 mosquito-borne pathogen species (Fig. 2c), often detected in several mosquito species, including across genera (Fig. 2c). The number of pathogen species detected per mosquito species increased with the mosquito species sample size, albeit in a non-monotonic way (Fig. S5). The mean mosquito species per pathogen species was 1 for filariae, 1.7 for arboviruses, and 3.9 for plasmodia. With nine species infected with at least one mosquito-borne pathogen species, *Culex* dominated this list (Fig. 2c and S4). Seven mosquito species exhibited disseminated infections for at least one pathogen (Fig. 2c). Pathogen richness in *Cx. perexiguus* was highest (11), including two arboviruses (WNV, MPOV) and 9 plasmodia (Fig. 2c and S4) compared with a mean of 2.9 mosquito-borne pathogens per mosquito species. It also had the highest richness of disseminated infections: six plasmodial species (only plasmodia exhibited head-thorax infections).

## Discussion

The old hypothesis that mosquito-borne pathogens are spread over large distances by windborne mosquitoes at altitude (Garrett-Jones 1950, 1956, 1962, Pedgley et al. 1995, Reynolds et al. 2006, Chapman et al. 2015, Huestis et al. 2019) was based only on epidemiological and meteorological data and on sporadic observations of mosquitoes at altitude (Glick 1939, Reynolds et al. 1996a). It was not widely accepted because it lacked direct evidence for i) the regularity and scales of such movements and ii) infection with mosquito-borne pathogens in windborne mosquitoes at altitude. Systematic aerial collections over Africa have revealed regular, large-scale windborne migration of mosquitoes at altitude (Huestis et al. 2019, Yaro et al. 2022, Atieli et al. 2023), and our results reveal high infection rates with arboviruses, plasmodia, and filariae (overall 12.7%). Additionally, high rates of disseminated infections (overall 6.3%) implicated such mosquitoes as probably infectious—ready to transmit pathogens when taking their first blood meal after landing (Sanogo et al. 2021). Although salivary gland infection barriers can render mosquitoes with disseminated infections unable to transmit, these barriers are rare (Hardy 1988). Overall, 21 mosquito species were found infected and 15 were infectious with flaviviruses, plasmodia, or filariae, indicating that windborne migration is widespread across pathogens exploiting multiple mosquito vectors species rather than concentrated in few super-infected mosquitoes (Fig. 2).

Remarkably, 19 mosquito-borne pathogen species were identified from this modest sample size (1,009 female mosquitoes), including arboviruses affecting humans (dengue, West Nile, and M’Poko viruses). Infection, even with macro-parasites such as filarial nematodes, clearly does not prevent mosquitoes from undertaking high-altitude flights as suggested previously (Garms et al. 1979, Garms 1981, Cheke et al. 2024). These results provide compelling evidence that mosquito-borne pathogens are often spread by windborne mosquitoes at altitude. Here, we describe, for the first time, the composition of mosquito-borne pathogens and their high-altitude mosquito carriers over Africa and discuss its implications.

The importance of windborne pathogen spread by mosquitoes relies on the species composition of the pathogens and the mosquitoes at altitude. Additionally, it relies on mosquito flight duration, the number of high-altitude journeys (nights) individual mosquito takes, selectivity of wind direction and speed, and whether these mosquitoes survive their high-altitude transport and refeed upon their descent. Hence, it also depends on the suitability of the landing destinations to the mosquitoes including abundance of susceptible hosts there, and mosquito age that predicts her remaining reproductive cycles and transmission events. If pathogens arrive to the same destinations using migratory birds (Moon et al. 2019), or human transport (Kilpatrick et al. 2006, Nartey et al. 2024), the importance of windborne mosquito spread diminishes.

The absence of human plasmodia, so common in people (65% in Mali, (Dao et al. 2023), 73% in Ghana (Heinemann et al. 2020)) suggests that the majority of these mosquitoes have fed on animals rather than people. Further, at least 18 of the 19 mosquito-borne pathogen species detected circulate among wild animals and are considered sylvatic, e.g., dengue virus also circulates among nonhuman primates and possibly birds (Gwee et al. 2021). This highlights the value of aerial collections in surveillance of sylvatic pathogens that are especially difficult to monitor; many even lack diagnostic assays (Eastwood et al. 2020, Figuerola et al. 2022, Lehmann et al. 2023a). Pathogen spread by windborne mosquitoes likely connects sylvatic foci and may prevent regional extirpation despite fluctuations including loss in some foci (Ericson et al. 1999, Laine 2005, Kerr et al. 2006, Harding et al. 2012), thus playing key role in maintenance of these pathogens. Although the importance of pathogen spread via windborne mosquitoes diminishes if they infect highly mobile hosts such as migratory birds that may be even more efficient at dispersing mosquito-borne pathogens, windborne mosquitoes likely arrive in some destinations that migratory birds avoid because they are situated away from their flyways (Moon et al. 2019) or because their step-length is too large.

Seventeen of 19 mosquito-borne pathogens we detected circulate in birds (Table 1). This probably reflects the predominance of species of *Culex* (75.1%) and *Coquillettidia* (5.9%) that commonly prefer feeding on birds (Soghigian et al. 2023). Although prevalence was not related to species sample size (Figures 2 and S2), the number of pathogens detected per mosquito species increased with sample size (Figure S4), providing possible explanation for the predominance of avian pathogens. Sampling larger numbers of mosquitoes that feed on mammals, such as *Aedes* and *Anopheles* will likely increase representation of mammalian pathogens. Evidence suggests that birds exhibit persistent infections with plasmodium (even over years) and exceptionally high prevalence (>90%) (van Rooyen et al. 2013). Birds exhibit less extreme but consistent trend with arboviruses (Wheeler et al. 2012). It is unlikely that feeding on birds increases the likelihood to engage in high-altitude migration, but this remains unclear.

Although *Cx. watti* was the dominant species in our aerial collection, *Cx. perexiguus* (the 2^nd^ most dominant) exhibited the highest number of pathogens per mosquito species as well as the number of pathogens with disseminated infections and it also showed a relatively high infection and infectiousness rates (Figures 2, S2, and S4). Multiple studies have established or implicated its role in the transmission of WNV, Usutu virus, Bagaza virus, Sindbis virus, and various plasmodia in Africa, Europe, and the Mediterranean e.g., (Ayhan et al. 2022, Figuerola et al. 2022). Because of its large geographical range across Africa, southern Europe, and Asia (Lehmann et al. 2023), its ranking as the second most abundant mosquito species at altitude in our collection (Fig. 1) and the most abundant over the Sahel (Yaro et al. 2022), this species is flagged as a particularly influential species on windborne pathogen spread.

Given that many mosquito species bloodfeed preferentially on multiple vertebrate species (Tedrow et al. 2019, Soghigian et al. 2023), we expect that the 61 species, representing 9 mosquito genera (Fig. 1a) feed on diverse set of vertebrate species, and therefore are likely to spread diverse mosquito-borne pathogens (Tandina et al. 2018, Eastwood et al. 2020, Lehmann et al. 2023a). Because the combined mosquito faunas in Mali (105 species) and Ghana (155 species) comprises 181 species (Lehmann et al. 2023a), our aerial collection sampled 34% (61/181) of this fauna. This fraction is likely an underestimate given the modest sample size taken from five stations in an ecologically diverse region of approximately 1,500,000 km^2^. Estimates of the species’ aerial densities reveal that a panel density of a single mosquito throughout the study (191 collection nights), is equivalent to an aerial density of 1 female mosquito/390 million m^3^ of air. Hence, the corresponding number of mosquitoes expected to cross a 100 km line perpendicular to the wind direction between 100 and 300 m agl is 11,635 per night. Accordingly, during a migration season of 4 months, >one million mosquitoes of each of the 21 rarest species fly at altitude over that sector alone, while 10—200 million for each of the 12 most common species. Conservatively using a disseminated infection rate of 0.1% (0.1—4.6%, Table 1) implies thousands of infectious high-altitude windborne mosquitoes per species crossing each 100 km sector. Gravid mosquitoes survive >11 hours at altitude, readily lay eggs afterwards and take a new bloodmeal (Sanogo et al. 2020, Atieli et al. 2023). Therefore, the likelihood of pathogen spread over tens or hundreds of kilometers away from endemic area or sylvatic foci cannot be disregarded (Huestis et al. 2019, Yaro et al. 2022).

## Acknowledgements

We thank Drs. Jose’ Ribeiro, Thomas Wellems, Carolina Barillas-Mury, Jesus Valenzuela, Thomas Mrs. Fatoumata Bathily, and Mrs. Margery Sullivan for their kind and vital support. Drs. Alvaro Molina-Cruz and Fabiano Oliveira for their valuable guidance and assistance with metagenomics and molecular analyses. Thanks to Ms. Mona Kafaie for assistance in laboratory work. Special thanks to the residents of the villages Thierola, Kenieroba, Bia, Dukusen, and Wenchi for their cooperation and hospitality. This study was supported by the Division of Intramural Research, National Institute of Allergy and Infectious Diseases, National Institutes of Health, Bethesda MD (ZIA AI001196-06), the Bill & Melinda Gates Foundation, Grand Challenges Explorations grant (OPP1217659; awarded to TL), the West African Center for Emerging Infectious Disease (NIH/NIAID grant U01 AI151801).

## Materials and Methods

### Study sites

Aerial collection stations were established in open rural areas in Mali and Ghana: the Sahelian village, Thierola (13.659, -7.215, Mali), the Sudano-savanna village, Kenieroba (12.112, -8.332, Mali), the Guinea-savanna village, Bia (10.492, -5.910, Mali), the Guinea woodland ecozone near the town of Wenchi (7.781, -2.162, Ghana), and a moist-semi-deciduous forest near the town of Agogo (6.961, -0.960, Ghana). Sites from Mali received annual precipitation of 500—900 mm during the short-wet season (June-October) whereas in Ghana, annual precipitation varies between 1,200—1,500 mm, and spread during most months of the year (Siebert 2014). These study areas have been described previously (Dao et al. 2014, Huestis et al. 2019, Nartey et al. 2024), as have the field methods used in this study (Huestis *et al*., 2019; Florio *et al*., 2020; Sanogo *et al*., 2021). Collections were made between March 2018 and October 2022. Due to logistical reasons sampling intensity differed between localities with 50, 16, 67, 26, and 32 sampling nights in Thierola, Kenieroba, Bia, Wenchi, and Agogo, respectively.

### Aerial collection

The methods have been described in detail previously (Huestis et al. 2019, Florio et al. 2020, Sanogo et al. 2021). Briefly, insect sampling was conducted using sticky nets (1 x 3 m panels) attached to the tethering line of helium-filled balloon (3 m or 3.3 m diameter), with each balloon typically carrying three panels. Panels were suspended at 120 m, and 160 m 190 m agl on the 3 m balloon and 120, 190 m, 240 m, and 290 m agl on the 3.3 m balloon. Balloons were launched approximately one hour before sunset and retrieved one hour after sunrise, the following morning. To control for insects trapped near the ground as the panels were raised and lowered, comparable control panels were raised up to 100 m agl and immediately retrieved during each balloon launch and retrieval operation. Following panel retrieval, inspection for mosquitoes was typically carried out immediately and specimens removed from the nets with forceps, were briefly washed in chloroform to remove the insect-glue and individually stored in labeled vials containing RNAlater™ Stabilization Solution (Invitrogen, Thermo Fisher Scientific, US). After several days in room temperatures in field conditions, specimens were placed in -20C freezers. Other insects were stored in vials containing 80% ethanol.

### Specimen processing and DNA/RNA extraction

In the laboratory, mosquitoes were thawed over ice, placed momentarily on filter paper to absorb excess RNAlater^TM^ solution, washed in deionized water and examined under dissecting microscope. Specimens were identified to genus or subfamily, their sex and gonotrophic state observed and recorded. Initially, DNA/RNA extraction was carried on whole specimens. Later, the female’s abdomen was dissected from her thorax and each part was independently subjected to nucleic acids extraction as were whole bodies of male mosquitoes. Extractions were carried out using Trizol (TRI Reagent®, Zymo Research, US) after tissue homogenization using low-bind beads (in a Mini-BeadBeater-96 (BioSpec Products, Inc., Bartlesville, OK, USA) at a max speed for 30 s, repeated three times with 30 sec intermissions to dissipate heat. This slurry was spun at 13,000 g at 4C for 5 min to clear solids, and the supernatant was used for the extraction using Mag-Bind® Viral DNA/RNA 96 Kit (Zymo Research, US) using the Kingfisher Robot (KingFisher® Flex, ThermoFisher, US). The extracted nucleic acids were suspended in 50 µl of molecular grade water. No DNAses were used to preserve both the RNA and the DNA.

### Mosquito identification and pathogen screening

The mosquito mitochondrial COI gene was PCR amplified (Folmer et al 1994) using barcoded primers (identifying each amplicon) and 1.5 µl of the nucleic acids extracts (above). The 658 bp amplicons were sequenced using the Oxford Nanopore technologies long-read MinION NGS platform, following established protocols (Srivathsan et al. 2019, Srivathsan et al. 2024). A small minority of the specimens were subjected to standard PCR followed by Sanger sequencing (Eurofins Genomics, USA) of their amplicons after purification using QIAquick PCR Purification Kit (Qiagen, USA).

Samples consisting of RNA/DNA extractions of females were screened individually for pathogen groups including flaviviruses, alphaviruses, plasmodia, and filariae using group-specific real-time PCR assays with primers (and associated probes, Table S5). To detect infection with plasmodia, mosquitoes were screened with Plasmodium-genus qPCR targeting the *cytochrome oxidase gene-1* (*COI*) following (Mediannikov et al. 2013). Positive samples, defined as having CT<36 (Diarra et al. 2017) were subjected to nested PCR amplification targeting 477 bp and 799 bp of *cytochrome b* (*cyt b*) (Hellgreen et al. 2004; Pekins and Schalk, 2002; Templeton et al 2016) (Table S5). Anopheline mosquitoes were also screened for human plasmodia using the qPCR assays of Bass et al. (2008).

To detect infection with filarial nematodes, mosquito samples were screen using qPCR targeting *28S rRNA gene* (Laidoudi et al. 2020). Positive samples were subjected to standard PCR of the *cox gene* were confirmed by visualizing amplicons on 2% agarose gel (Laidoudi et al. 2020) and sequenced using Sanger sequencing (Eurofins Genomic, USA). Infection with arboviruses of the genera Orthoflavivirus (and Alphavirus) were screened in the abdomens and head-thorax sections of mosquitoes by RT-qPCR associated with high-resolution melting curve (Vina-Rodriguez et al 2017) using one step mixes (GoTaq® 1-Step RT-qPCR System, Promega, US). Positive flaviviruses samples were subjected to a nested PCR on a 960 bp fragment of the NsP5 gene following Vazquez et al. (2012), and purified amplicons were sequenced by Sanger sequencing (Eurofins Genomics, MD US). A subset of 48 samples were sequenced by MinION Oxford nanopore (Below).

All PCR and qPCR assays had at least 1 negative and 1 positive controls per assay. Molecular grade water was used as negative control and corresponding sample (e.g., cultured *P. falciparum*, *Brugia malayi*/*W. bancrofti*, dengue virus, and Eilat virus were used as positive controls for the pan-*Plasmodium*, pan-nematode, pan-flavivirus, and pan-alphavirus assays, respectively. In the case of dengue, the ∼960 bp long sequences of the NS gene of the two wild infected mosquitoes differed from the positive control by 5 and 9 substitutions. The result was independently repeated for verification.

### Metagenomics on selected positive samples

Metagenomic analyses on 48 individual mosquito abdomens found to be positive for flaviviruses (N=29) or plasmodium (n=10) or filaria (N=9) divided into two pools of 24 samples were carried out using MinIon nanopore platform in two separate flow cells following methods as previously described (Ergunay et al. 2023). Briefly, the cDNA library was prepared as previously described (Kamau et al. 2022). Briefly, using the RevertAid First Strand cDNA Synthesis Kit (Thermo Fisher Scientific), purified nucleic acids were converted to complementary DNA (cDNA) while random primer mix (New England Biolabs, Ipswich, MA, USA) was used to prepare double-stranded cDNA using NEBNext Non-Directional RNA Second Strand Synthesis module. Barcoded cDNA with the Native Barcoding Expansion 96-EXP-NBD 196 (Oxford Nanopore Technologies, Oxford, UK) was synthesized using automatic device, epiMotion 5075 (Eppendorf, US). Double stranded cDNA was cleaned up using Agencourt AMPure XP reagent (Beckman Coulter Biosciences, Indianapolis, IN, USA) and quantitated by Qubit dsDNA HS Assay Kit (ThermoFisher Scientific, Waltham, MA, USA). The NEBNext Ultra II End repair/dA-tailing and Quick Ligation modules (New England Biolabs, Ipswich, MA, USA), and Ligation Sequencing Kit SQK-LSK114 (Oxford Nanopore Technologies, Oxford, UK) were used to prepare libraries, which were further quantitated by Qubit (ThermoFisher Scientific, Waltham, MA, USA) using 1 × dsDNA HS Assay Kit. Metagenomic sequencing was accomplished using one pool (24 samples) per flow cell (Flow Cell (R10.4.1), and run on the GridION sequencing device (Oxford Nanopore Technologies, Oxford, UK) for 72 h.

Following nanopore discovery-metagenomic sequencing, base-calling and demultiplexing was accomplished on the device with the MinKNOW operating software v21.11.7 (Oxford Nanopore Technologies) and Guppy v5.1.13 (Wick et al. 2019). Raw reads were trimmed with Porechop to remove adapter sequences and then filtered with NanoFilt to remove reads with q-scores ≤ 9 and read lengths ≤ 100bp (Wick et al. 2019). Host mosquito genomes were further removed using Minimap2 v2.24 and Samtools v1.9 (Danecek et al., 2021; Li et al., 2018). Subsequently, the data was aligned to the National Center for Biotechnology Information (NCBI) non-redundant (NR) database using DIAMOND v2.0.14 and visualized using MEGAN6 (v6.25.9)(Huson et al. 2016)). Sequences were handled using Geneious Prime (v2023.2.1) (Biomatters Ltd., Auckland, New Zealand). De novo assembly and mapping were carried out using Geneious Prime plug-ins, with default settings optimized for nanopore data. BLASTn and BLASTp algorithms were used for similarity searches in the NCBI database. CLUSTALW was used for sequence alignment and pairwise comparisons (Thompson et al., 1994).

### Data analysis

Data from the mosquito mtCOI amplicon sequencing generated using the minION were analyzed through ONTbarcoder version 1.9 or 2.0. Sequences of individual mosquitoes (amplicon metagenomics consensus per individual) and pathogens obtained by Sanger sequencing were blasted against repositories in BOLD, GenBank (NCBI) and, unless otherwise specified, species identity was provisionally given if sequence similarity was ≥98% (in most cases it was ≥99%). Neighbor joining (NJ) phylogenetic trees were used to cluster the sequences derived here in comparison to best matches from corresponding databases. Mosquito taxa that did not fit these criteria were provisionally named (e.g., *Aedes mali* sp. 1*, Culex* MBI-61) based on their sequence (or subfamily), in cases where amplification failed repeatedly, but the specimen was visually identified consistently in the field and in the laboratory, they were identified by subfamily: Culicine or Anopheline.

Phylogenetic relationships among 78 *Plasmodium* partial *cytb* gene sequences (470 bp) obtained in this study and previously reported haemosporidian sequences were estimated on an alignment performed using ClustalX v2.0.12 and Muscle as implemented in SeaView v4.3.5 (Gouy et al. 2010) with manual editing. Phylogenetic hypotheses were assessed based on this alignment using a Bayesian method implemented in MrBayes v3.2.7 with the default priors (Ronquist and Huelsenbeck 2003), and a general time-reversible model with gamma-distributed substitution rates, and a proportion of invariant sites (GTR + Γ + I) as it was the best model that fit the data with the lowest Bayesian information criterion scores estimated by MEGA v7.0.26 (Kumar et al. 2016). Bayesian supports were inferred for the nodes in MrBayes by sampling every 1000 generations from 2 independent chains lasting 2 × 106 Markov Chain Monte Carlo steps. The chains were assumed to have converged once the potential scale reduction factor value was between 1.00 and 1.02, and the average standard deviation of the posterior probability was <0.01. 25% of the samples were discarded as a ‘burn-in’ once convergence was reached. Lineages names of all sequences (partial *cytb* gene) used here are shown in the phylogenetic trees and new isolates were named after their mosquito identification code. The phylogenetic tree of the plasmodia isolates was used to confirm species assignment by comparison to sequences of taxa included in the MalAvi (Bensch et al. 2009) and NCBI databases. The latter comprised of 40 published sequences including *P. matutinum*, *P. vaughani*, *P. relictum*, *P. cathemerium* as well as the closely related sister taxa *Haemoproteus coraciae* and *Leucocytozoon* sp. as outgroups to root and stabilize the tree topology (supp. Fig. 2). Plasmodial taxa that mapped >3% sequence divergence from their nearest neighbor and away from their named best match were provisionally named *P. mali sp. 1*, etc.

Species-specific mosquito exposure to blood was estimated as the fraction of gravid, semi-gravid, and blood fed females combined as opposed to unfeds. Species-specific and whole sample infection rates were estimated across mosquito specimens with data on particular pathogen (e.g., DENV) and groups of pathogens (flaviviruses, or vertebrate pathogens). Overall mosquito infection rate was estimated based on detection of either flavivirus, plasmodia, or nematode in any part of a mosquito (data on alphaviruses were not used because sequencing positives failed to confirm infection detected by the qPCR assay). Mosquitoes that were infected with more than one pathogen, or in both body parts were given the same weight as if they were infected with one. Infectiousness or disseminated infections were calculated based on the thorax and head body part alone. Abdominal infection rates were estimated based that body part alone.

#### Statistical analysis

Heterogeneity among species in sex, gonotrophic state, and infection compositions were evaluated using contingency table likelihood ratio chi-square test after pooling species with sample size<6. If the fraction of cells with expected counts <5 were greater than 20%, we used exact tests based on Monte Carlo simulation of 10,000 samples using Proc Freq (SAS Institute, 2012). Weighted regression analysis (using sample size per species as weight) and local regression to assess trends between variables were calculated using Proc Reg statistical graphics facilities (Proc Sgplot) in SAS (SAS Institute, 2012).

Aerial density was estimated using the panel density of the species divided by the total air volume that passed through that net that night (i.e., aerial density = panel density/volume of air sampled, and volume of air sampled = panel surface area × mean nightly wind speed × sampling duration). The panel surface area was 3 m^2^. Wind-speed data were obtained from the atmospheric re-analyses of the global climate (ERA5). Hourly data consistent of the eastward and northward components (horizontal vectors) of the wind were available at 31-km surface resolution at 2 and 300 m agl (1,000 and 975 mbar pressure levels). Overnight records (19:00 through to 06:00) were averaged to calculate the nightly mean direction and mean wind speed over each African sampling station based on standard formulae using code written in Base SAS (SAS Institute, 2012). Nightly collections reflecting duration panels were suspended at altitude lasted 14 hours. The intensity of migration was expressed as the expected number of migrants/species crossing a line of 1 km perpendicular to the wind direction at altitude, which reflect their direction of movement (Drake and Reynolds 2012; Hu et al. 2016; Reynolds et al. 2017; Florio et al. 2020). We used the mean wind speed at altitude during the migration season (4.5 m/s) and assumed that the mosquitoes fly in a layer depth of 200 m agl (Huestis et al. 2019, Florio et al. 2020). The nightly migration intensity was computed across the four months flight season (in which most species were sampled) including sampling nights during which no migrants were captured). The corresponding annual index was estimated for a sector of 100 km following (Huestis et al. 2019, Florio et al. 2020).

## Supplemental Results & Discussion

In this study we included samples collected at five aerial stations across West Africa to address the question of infection of high-altitude windborne mosquitoes with pathogens. A total of 1,249 mosquitoes were collected on 432 standard panels but no mosquito was collected on 301 control panels, demonstrating that mosquitoes were collected at altitude. Larger number female mosquitoes was sampled in Ghana (656; Agogo and Wenchi) than in Mali (361; Bia, Kenieroba, and Thierola, Fig. S1). A comparison of the high-altitude mosquito and pathogen compositions among these sites is beyond the scope of the present study and will require a larger sample size from most sites. The results revealed that mosquitoes infected with mosquito-borne pathogens were found in all sites (not shown).

Mosquito composition in altitude (Fig. 1a, Table S1) is typical of biological samples in being dominated by few common species whilst most species are represented by moderately frequent species and by rare species (Anderwartha and Birch 1954). This distribution is probably a reflection of both the abundance of these species on the ground over large catchment area as well as their aptitude to engage in high altitude flight. Although most rare species are less abundant at altitude, some may represent incidental events. For example, the two *Ae. aegypti*, a diurnal species that are attracted to people, were collected on the same panel during the first month of operation of a newly trained team. Additional studies will ascertain the status of rare species.

Consistent with previous studies (Huestis et al. 2019, Yaro et al. 2022, Atieli et al. 2023), at altitude female mosquitoes predominated (85%) reflecting a general species- and genera- independent pattern, because no heterogeneity was detected (Table S2). This asymmetry likely reflects sexual fitness differential associated with long-range migration. Accordingly, upon landing females are likely to find suitable larval sites with equal or higher prospects for larval development success, whereas males need to locate virgin females where they might be scarce and compete with local males that are expected to be in similar density to females—same ratio as in their provenance site. Although, mating in a new locality is expected to also increase male fitness, but the expected fitness increase is smaller due to the additional costs of locating virgin females and successfully competing with local males. Because females are probably inseminated by males from their provenance site, this asymmetry in long-range migration should not have consequences on sex-specific geneflow (Barth et al. 2013).

Heterogeneity in the fraction of gravid and unfed females has been detected among species (Table S3). Because all gravid females had at least one exposure to vertebrate blood whereas some of the unfed females may have not had such exposure, it is of interest *Cx. perexiguus* exhibits the highest fraction of gravid females (65%), which could further contribute for its high infection rate with diverse pathogens. (main text).

Younger mosquitoes have longer remaining life span, but older mosquitoes are more likely to be infected. The ratio of infectiousness/exposed (here, “infected” is inclusive for all body parts) mosquitoes can be used to estimate the fraction of mosquitoes above the minimum age to develop a disseminated infection (Gillies 1954, Gillies and Wilkes 1965). In tropical areas plasmodia typically require ≥7 days post infection to mature their oocysts in the midgut wall (abdomen) and release sporozoites into the haemocoel that accumulate in the salivary glands (thorax), thus initiating a disseminated infection. The first blood meal typically takes place on the adult’s second day (Gillies 1954, Gillies and Wilkes 1963, 1965), hence this ratio across all plasmodial species with at least 1 disseminated infection was 0.5 (0.031/0.062; Table 1), implying that half the population is ≥9 days old. This is 2.5 times the typical fraction of older mosquitoes that had taken ≥3 bloodmeals on vertebrate hosts (Gillies and Wilkes 1965), thus explaining, at least in part, the high infection rates in this collection. Whether pathogens increase aptitude of infected mosquitoes to engage in high-altitude windborne migration remains to be answered (Lion et al. 2006, Martini et al. 2015, Poulin and Dutra 2021). Novel methods to track mosquitoes over large distances and methods to determine their provenance would help address key questions on these journeys.

Finally, we touch on the question, how was the spread of pathogens by mosquitoes at altitude not already widely recognized by epidemiologists? The high mobility and near-continuous distribution of human and domestic animals led to the assumption that they are the primary long-range movers of their pathogens including mosquito-borne pathogens, alternatives were disregarded without evaluation. Inferring windborne-mosquito spread was deemed possible only when outbreaks occurred outside the disease typical range, vertebrate movement was soundly ruled out, and wind direction and speed aligned with the nearest known source (Garrett-Jones 1950, 1956, 1962, Sellers et al. 1977, 1982, Pedgley 1983, Sellers and Maarouf 1990); conditions that greatly limit discovery of pathogen spread by windborne mosquitoes even if it was the most common mode of spread. Additionally, this epidemiological analysis was applied primarily to zoonotic pathogens affecting people and domestic animals, thus missing most sylvatic pathogens that circulate among wild animals and represent the largest share of mosquito-borne pathogens in Africa (Lehmann et al. 2023). Infection rates for flaviviruses, Plasmodia, and filariae were higher in abdominal tissues than in head/thorax dissections (Fig. 1b, Tables 1 and S1).

---

**Figure S1.**
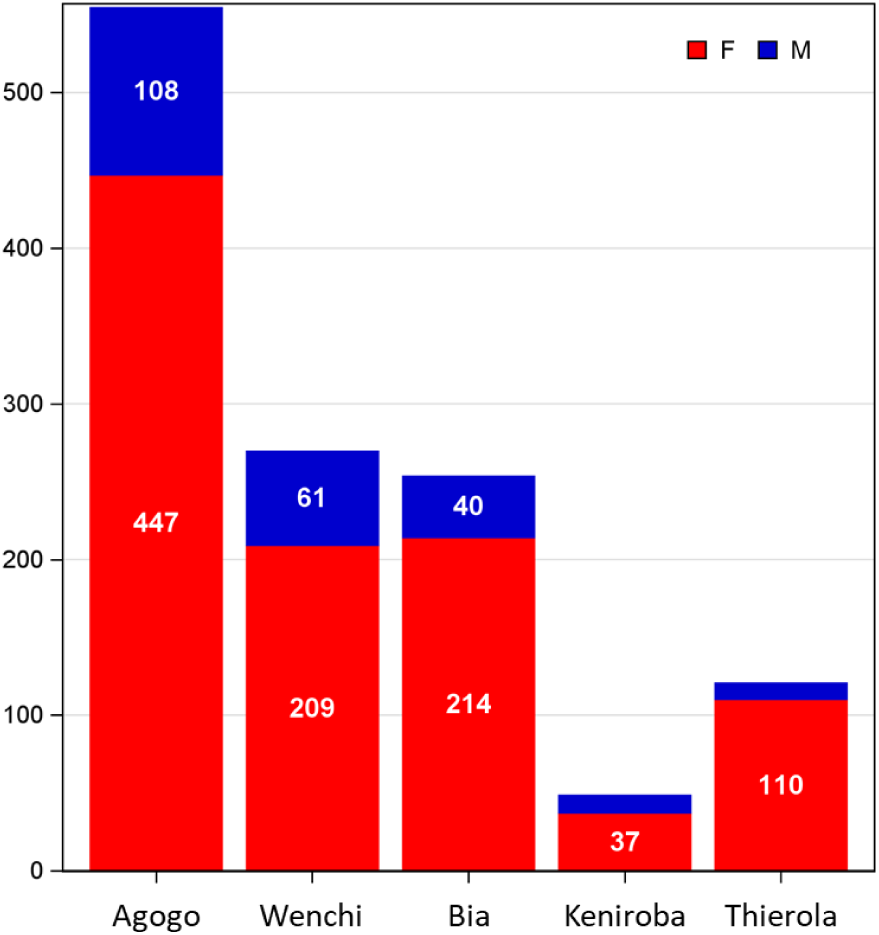
Sources of mosquitoes used in the present study among the aerial sampling stations by sex.

**Figure S2.**
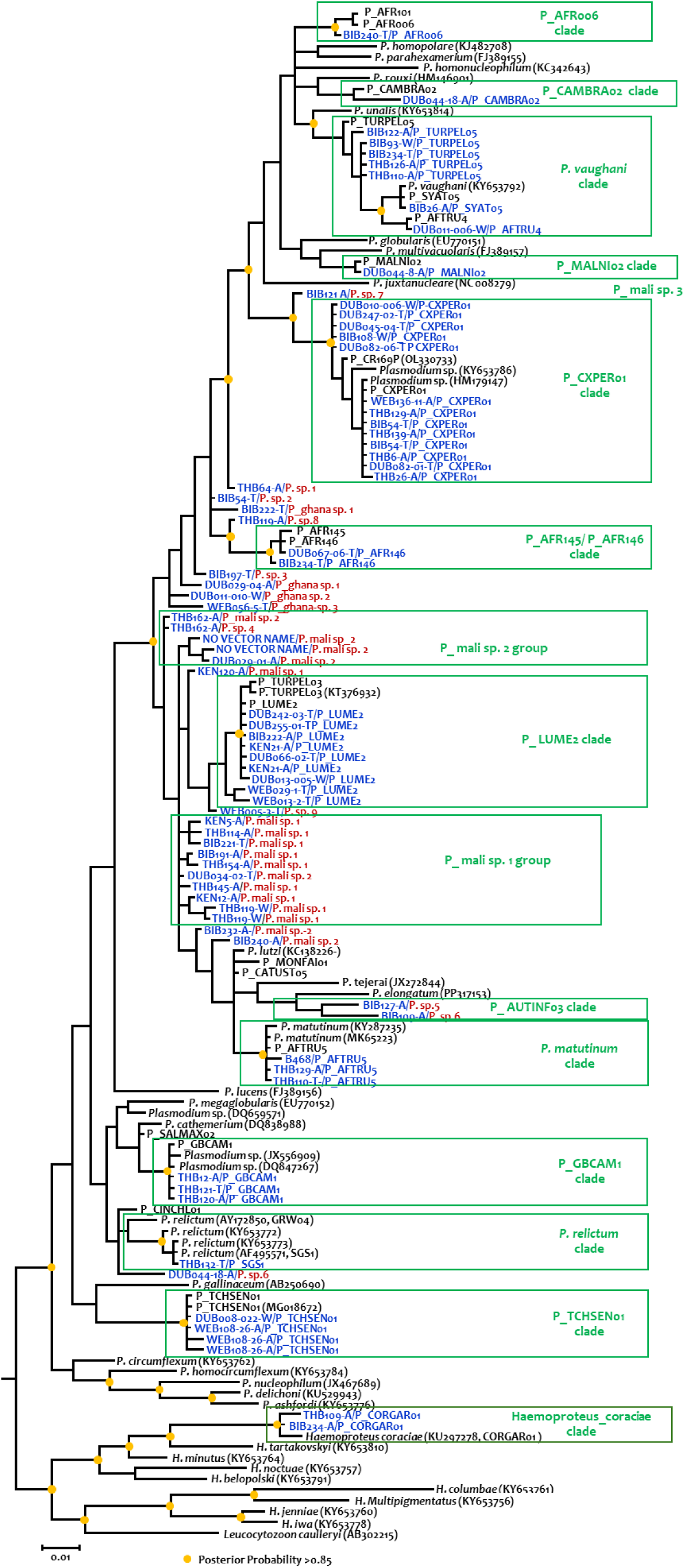
Bayesian phylogeny based on the mitochondrial *cytb* gene of plasmodia isolated from high-altitude mosquitoes (blue) compared with best-match sequences from MalAvi and NCBI databases (black). Posterior probabilities >0.85 are shown using a yellow circle. We grouped isolates separated by ≤3% sequence dissimilarity from closest neighbor to putative species and provisionally named them after the closest species or lineage in MalAvi that met this criterion (green boxes and fonts). Putative species that had no best match at this level were named sequentially, e.g., *P. mali sp. 1* after the country they were collected from. Note: *Haemoproteus* is closely related to *Plasmodium* but is vectored by non-mosquitoes biting flies.

**Figure S3.**
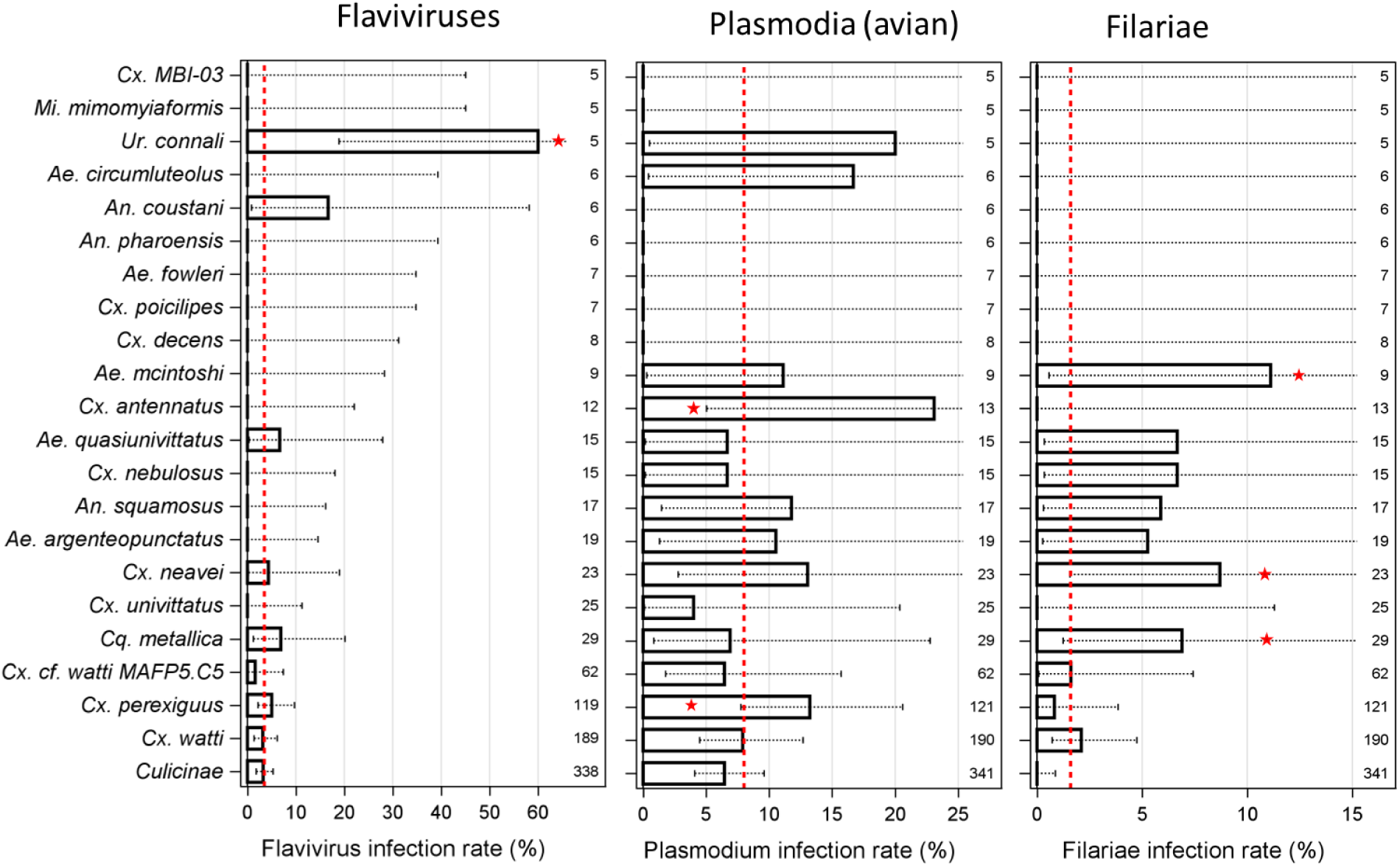
Overall infection rates per mosquito species (N≥5) based on pan-genus infection assays for flaviviruses (a), plasmodia (b), and filariae (c) with 90% CI. Higher infection rate (P<0.05, 2 side Binomial test) than the rate across all mosquito species (red lines) are indicated by stars.

**Figure S4.**
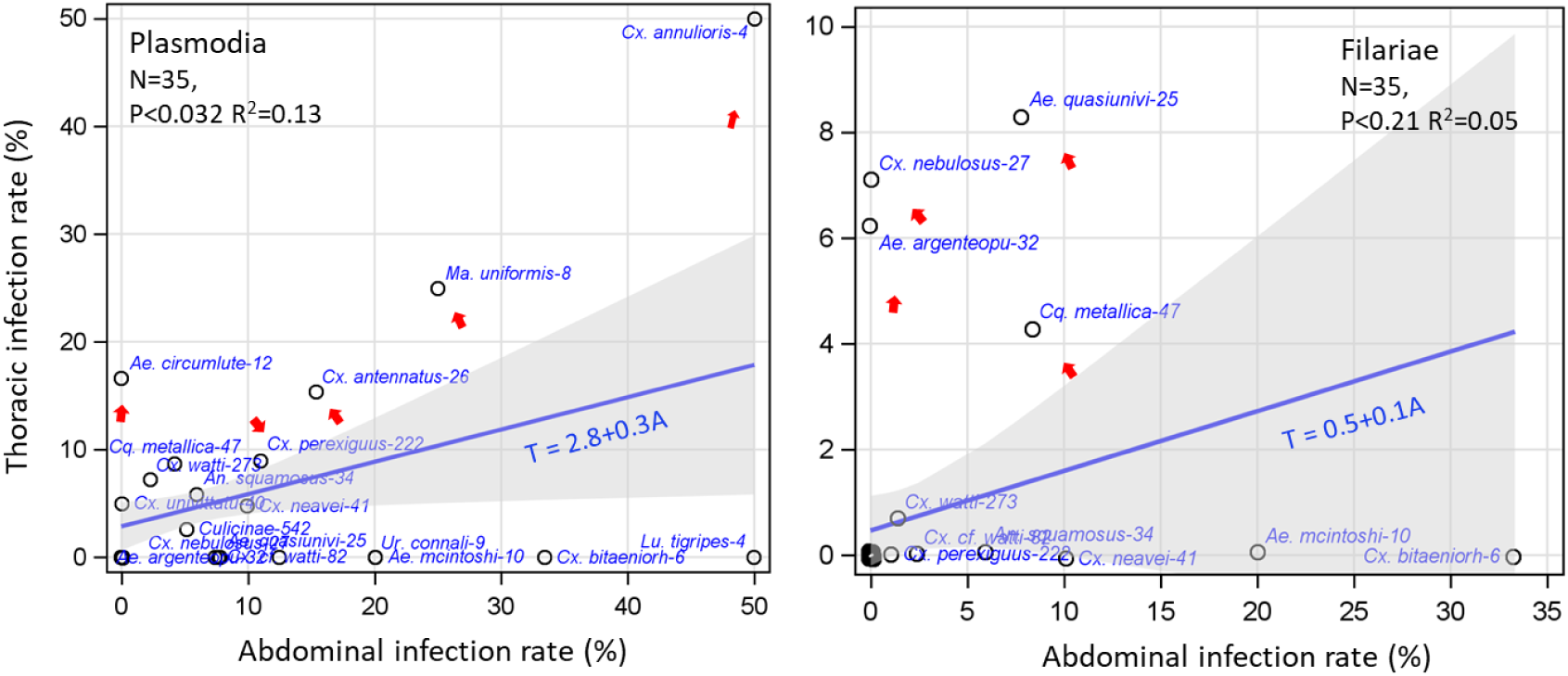
Relationship between disseminated (head-thorax) infection and exposure (abdominal) infection with plasmodia (a) and filariae (b) by species (abbreviated species name is listed with their sample size for infected species). Weighted linear regression (weighted by total sample size, blue) and 95% CI (gray) computed for species with N≥3 to identify outliers (red arrows) indicating competent vectors that typically feed preferentially on natural hosts, showing increased exposure.

**Figure S5.**
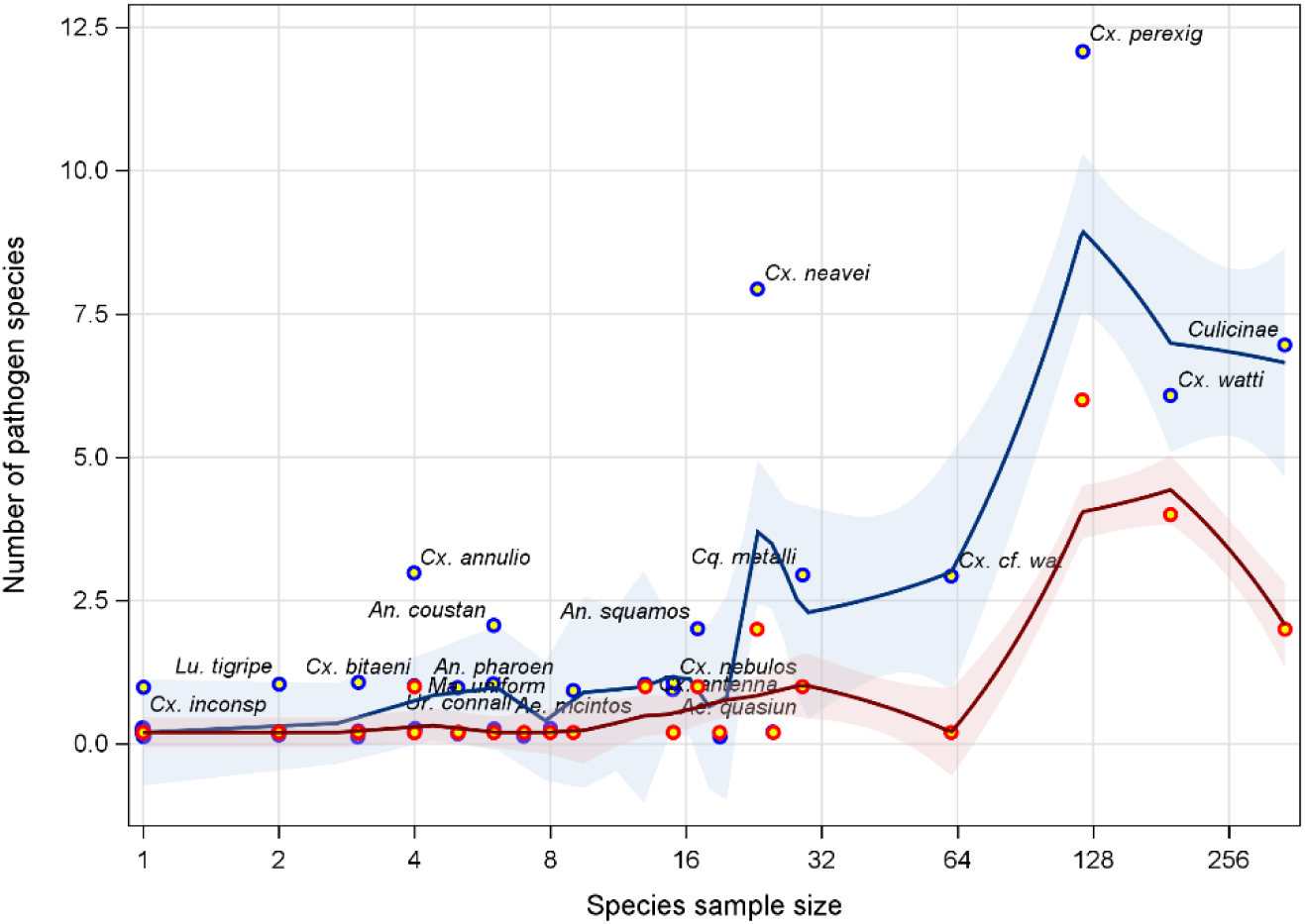
Effect of the species sample size on the number of pathogen species it is infected and infectious with. Loess regression lines with 95% CI describe the change in the number of pathogens species per mosquito species according to its ample size (log scale). Disseminated (head-thorax) infection (red) and total infection (blue) are shown. Abbreviated species name is listed for species with one or more pathogen species (black).

**Supplementary Table 1.** Composition of mosquitoes collected at altitude (120-290 m above ground; see main text)

| Species | COUNT | Comments |
| --- | --- | --- |
| *Ae. aegypti* | 2 | May represent contamination during panel processing: both specimens were collected on the same panel at the 1st month of operation of a new team |
| *Ae. argenteopunctatus* | 19 |  |
| *Ae. bromeliae* | 2 |  |
| *Ae. circumluteolus* | 8 |  |
| *Ae. dentatus* | 1 |  |
| *Ae. fowleri* | 7 |  |
| *Ae. mcintoshi* | 10 |  |
| *Ae. quasiunivittatus* | 18 |  |
| *Ae. sp. MBI-60* | 1 |  |
| *Ae. sp. MBI-61* | 1 |  |
| *Ae. sp. MBI-62* | 1 |  |
| *An. cf. rivulorum* | 1 |  |
| *An. coluzzii* | 2 |  |
| *An. coustani* | 7 |  |
| *An. funestus* | 1 | May represent contamination during panel processing: a single specimen that is recorded for the first time in altitude. Needs verification. |
| *An. gambiae s.l.* | 3 | May represent *Anopheles coluzzii* or *Anopheles gambiae* s.s. rather than a new species: molecular ID has failed repeatedly |
| *An. gambiae s.s.* | 1 |  |
| *An. pharoensis* | 6 |  |
| *An. pretoriensis* | 1 |  |
| *An. rufipes* | 3 |  |
| *An. squamosus* | 20 |  |
| *Anophelinae* | 2 | May contain additional species |
| *Cq. metallica* | 40 |  |
| *Cq. sp. MBI-01* | 5 |  |
| *Cq. sp. MBI-03* | 1 |  |
| *Cx. MBI-03* | 6 |  |
| *Cx. MBI-18* | 3 |  |
| *Cx. annulioris* | 5 |  |
| *Cx. antennatus* | 15 |  |
| *Cx. bitaeniorhynchus* | 4 |  |
| *Cx. cf. watti* MAFP5.C5 | 80 |  |
| *Cx. cinereus* | 1 |  |
| *Cx. decens* | 9 |  |
| *Cx. duttoni* | 1 |  |
| *Cx. ghana sp. 1* | 3 |  |
| *Cx. ghana sp. 2* | 3 |  |
| *Cx. inconspicuus* | 1 | May be a different species but closely related. To be checked again |
| *Cx. mali sp. 4* | 5 |  |
| *Cx. neavei* | 28 |  |
| *Cx. nebulosus* | 15 |  |
| *Cx. perexiguus* | 147 |  |
| *Cx. poicillipes* | 7 |  |
| *Cx. sp. MBI-60* | 1 |  |
| *Cx. sp. MBI-62* | 1 |  |
| *Cx. sp. MBI-65* | 1 |  |
| *Cx. sp. MBI-66* | 1 |  |
| *Cx. sp. MBI-68* | 1 |  |
| *Cx. sp. MBI-69* | 1 |  |
| *Cx. univittatus* | 32 |  |
| *Cx. univittatus B* | 4 |  |
| *Cx. watti* | 212 |  |
| *Er. sp. MBI-01* | 3 |  |
| *Lu. tigripes* | 3 |  |
| *Ma. africana* | 1 |  |
| *Ma. sp. MBI-60* | 1 |  |
| *Ma. uniformis* | 5 |  |
| *Mi. mediolineata* | 2 |  |
| *Mi. mimomyiaformis* | 5 |  |
| *Ur. alboabdominalis* | 5 |  |
| *Ur. connali* | 6 |  |
| *Ur. ghana sp. 1* | 3 |  |
| *Culicidae* (unidentified) | 5 | May contain additional species |
| *Culicinae* (unidentified) | 460 | Failed to produce PCR products or quality mtCOI sequence. May contain additional |
| Total mosquitoes intercepted | 1,249 | Few preserved in 80% ethanol or on silica gel (not in RNAlater) were not subjected to RNA |

**Table S2:** Sex composition by species in aerial collection (pooling species with N<10)

| spMo2 | Females % [N] |  |
| --- | --- | --- |
| Frequency<br>Percent<br>Row Pct | F | Total |
| **Ae. argenteopunctatus** | 19 | 19 |
|  | 2.43 | 2.43 |
|  | 100.00 |  |
| **Ae. mcintoshi** | 9 | 10 |
|  | 1.15 | 1.28 |
|  | 90.00 |  |
| **Ae. quasiunivittatus** | 16 | 18 |
|  | 2.05 | 2.30 |
|  | 88.89 |  |
| **An. squamosus** | 17 | 20 |
|  | 2.17 | 2.56 |
|  | 85.00 |  |
| **Cq. metallica** | 30 | 40 |
|  | 3.84 | 5.12 |
|  | 75.00 |  |
| **Cx. antennatus** | 13 | 15 |
|  | 1.66 | 1.92 |
|  | 86.67 |  |
| **Cx. cf. watti MAFP5.C5** | 65 | 80 |
|  | 8.31 | 10.23 |
|  | 81.25 |  |
| **Cx. neavei** | 23 | 28 |
|  | 2.94 | 3.58 |
|  | 82.14 |  |
| **Cx. nebulosus** | 15 | 15 |
|  | 1.92 | 1.92 |
|  | 100.00 |  |
| **Cx. perexiguus** | 122 | 147 |
|  | 15.60 | 18.80 |
|  | 82.99 |  |
| **Cx. univittatus** | 25 | 32 |
|  | 3.20 | 4.09 |
|  | 78.13 |  |
| **Cx. watti** | 190 | 212 |
|  | 24.30 | 27.11 |
|  | 89.62 |  |
| **PooSpecies** | 122 | 146 |
|  | 15.60 | 18.67 |
|  | 83.56 |  |
| **Total** | 666 | 782 |
|  | 85.17 | 100.00 |
| **Statistic** | **DF** | **Value; Prob** |
| **Chi-Square** | 12 | 16.2; 0.1817 |
| **Likelihood Ratio Chi-Square** | 12 | 20.9; 0.0522 |
| **Monte Carlo Estimate for the Exact Test** |  |  |
| **Pr >= ChiSq** | **0.0759** |  |
| **99% Lower Conf Limit** | 0.0691 |  |
| **99% Upper Conf Limit** | 0.0827 |  |
| **Sample Size** | 782 |  |
| **Number of Samples** | 10000 |  |

**Table S3:**
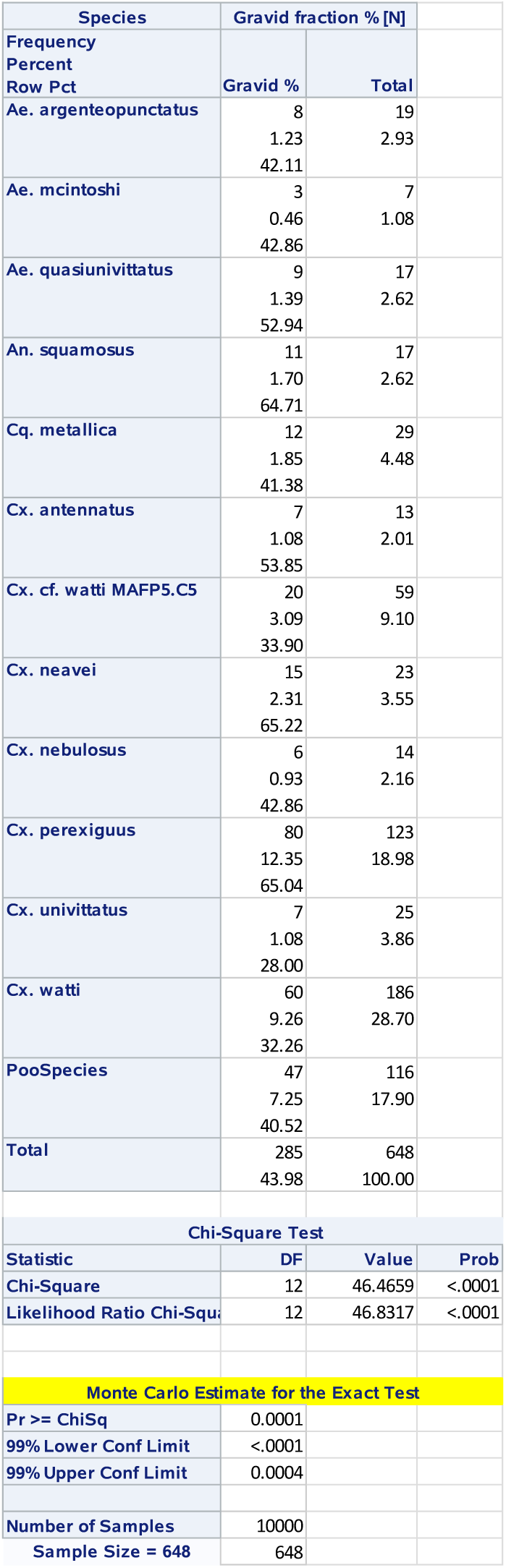
Fraction of gravid females in aerial collection by species (species with N<10 pooled)

**Table S4.**
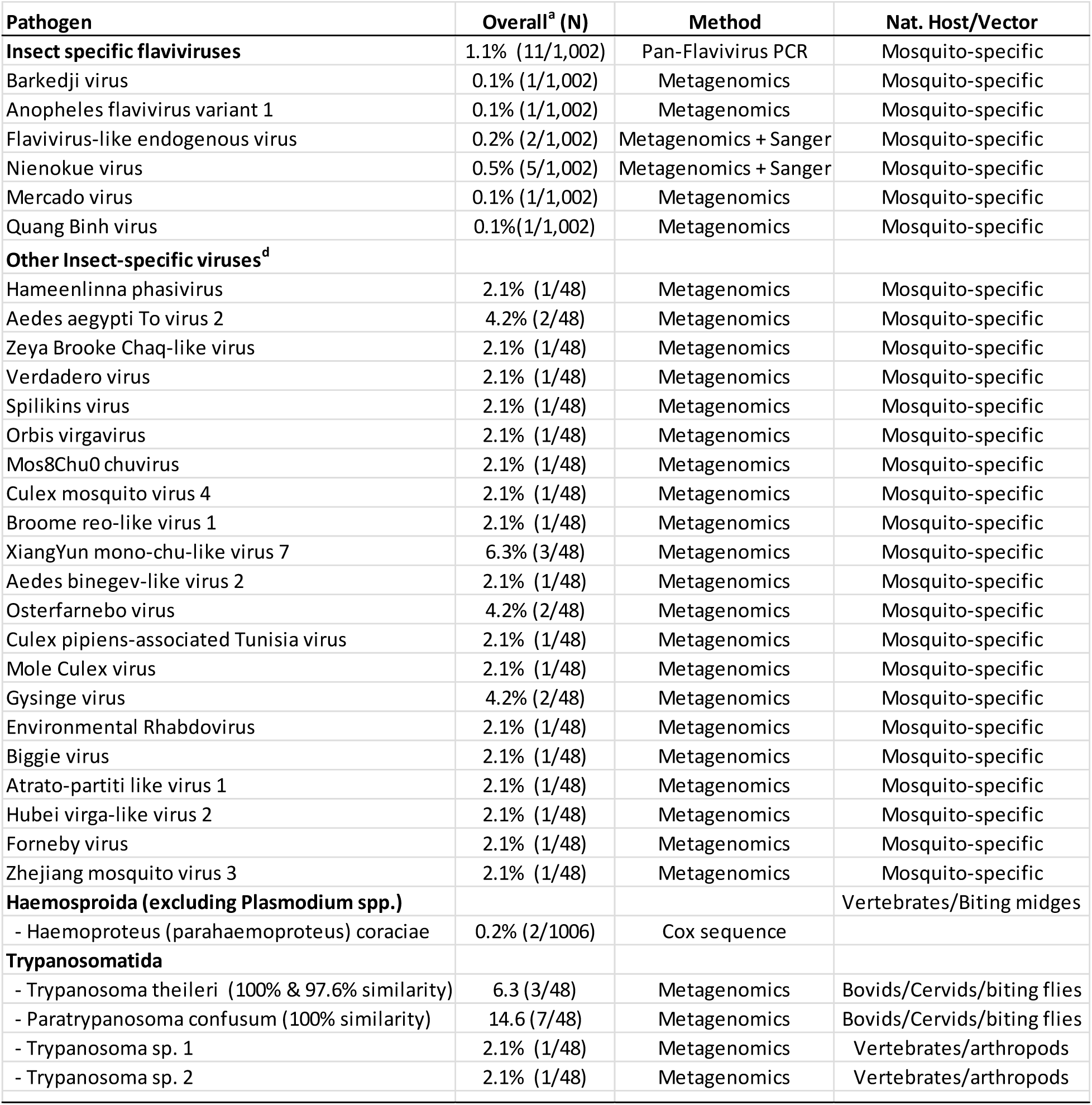
Infection rates of mosquitoes intercepted at altitude (120-290 m above ground) with insect-specific viruses and with non-mosquito-borne pathogens.

**Table S5.**
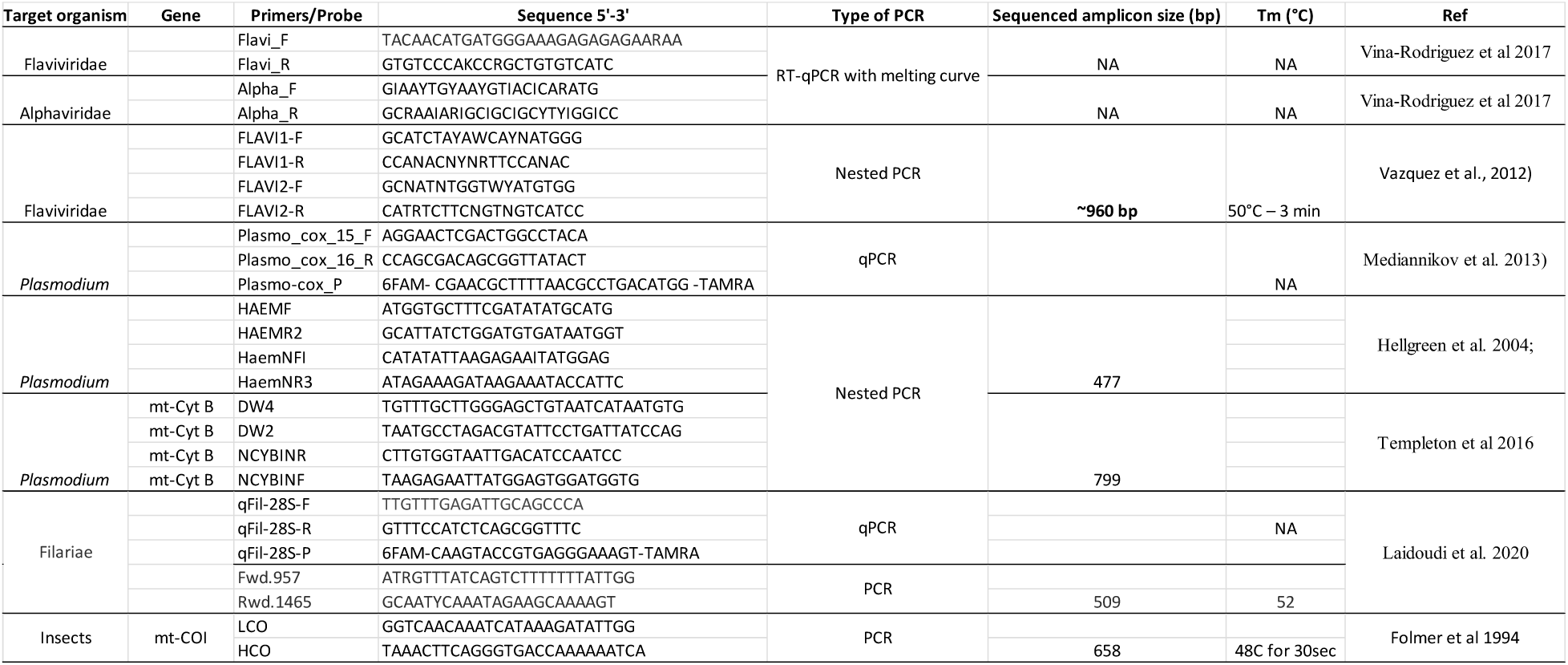
Primers used for pathogen and mosquito detection and identification.

## Notes

### Competing Interest Statement

The authors have declared no competing interest.

